# Enhancement of High-Density Lipoprotein-Associated Protease Inhibitor Activity Prevents Atherosclerosis Progression

**DOI:** 10.1101/2023.08.07.551670

**Authors:** Maura Mobilia, Alexander Karakashian, Callie Whitus, Khaga R. Neupane, Lance A. Johnson, Gregory A. Graf, Scott M. Gordon

**Author notes:** **Corresponding Author:** Scott M. Gordon Ph.D., Assistant Professor, University of Kentucky, Saha Cardiovascular Research Center, 741 South Limestone, BBSRB Room B349, Lexington, KY 40536-0509.

## Abstract

**Background:** Inflammatory cells within atherosclerotic lesions secrete various proteolytic enzymes that contribute to lesion progression and destabilization, increasing the risk for an acute cardiovascular event. The relative contributions of specific proteases to atherogenesis is not well understood. Elastase is a serine protease, secreted by macrophages and neutrophils, that may contribute to the development of unstable plaque. We have previously reported interaction of endogenous protease-inhibitor proteins with high-density lipoprotein (HDL), including alpha-1-antitrypsin, an inhibitor of elastase. These findings support a potential role for HDL as an endogenous modulator of protease activity. In this study, we test the hypothesis that enhancement of HDL-associated elastase inhibitor activity is protective against atherosclerotic lesion progression.

**Methods:** We designed an HDL-targeting protease inhibitor (HTPI) that binds to HDL and confers elastase inhibitor activity. Lipoprotein binding and the impact of HTPI on atherosclerosis was examined using mouse models.

**Results:** HTPI is a small (1.6 kDa) peptide with an elastase inhibitor domain, a soluble linker, and an HDL-targeting domain. When incubated with human plasma *ex vivo*, HTPI predominantly binds to HDL. Intravenous administration of HTPI to mice resulted in its binding to plasma HDL and increased elastase inhibitor activity on isolated HDL. Accumulation of HTPI within plaque was observed after systemic administration to *Apoe*^*-/-*^ mice. To examine the effect of HTPI treatment on atherosclerosis, prevention and progression studies were performed using *Ldlr*^*-/-*^ mice fed Western diet. In both study designs, HTPI-treated mice had reduced lipid deposition in plaque. Histology and immunofluorescence staining of aortic root sections were used to examine the impact of HTPI on lesion morphology and inflammatory features.

**Conclusions:** These data support the hypothesis that HDL-associated anti-elastase activity can improve the atheroprotective potential of HDL and highlight the potential utility of HDL enrichment with anti-protease activity as an approach for stabilization of atherosclerotic lesions.

## Introduction

Atherosclerosis is the underlying process behind coronary artery disease and the culprit for most acute cardiovascular events (1). Lipoproteins play key roles in both the development of and protection against atherosclerotic cardiovascular disease. Elevated plasma concentrations of low-density lipoprotein (LDL) contribute to atherogenesis (2). LDL particles infiltrate the vascular wall where they become trapped and modified by oxidative processes. Uptake of modified LDL by resident macrophages results in foam cell formation and the recruitment of additional inflammatory cells to the lesion, including neutrophils and monocyte derived macrophages (3-5). These cells secrete factors including cytokines and proteases, which can promote progression of the lesion, leading to larger plaque volume and a less stable morphology that is more prone to endothelial disruption and plaque rupture (6, 7). Neutrophil elastase (NE) is a serine protease that is abundant in atherosclerotic lesions and is secreted by both neutrophils and macrophages within the lesion (8). Proteolytic activity within lesions is thought to increase the propensity for atherothrombosis and the risk of an acute cardiovascular event such as myocardial infarction.

High density lipoproteins (HDL) are considered to be protective against atherosclerosis. The primary mechanism behind HDL-mediated atheroprotection has traditionally been attributed to HDL’s participation in cholesterol efflux and reverse cholesterol transport (9, 10). However, recent studies of the compositional complexity of HDL reveal that these lipoproteins may have other, more diverse roles in atherosclerosis. For example, HDL carry an extensive protein cargo that is enriched in proteins with roles in the regulation of protease activity (11, 12). Proteomics studies demonstrate that HDL associate with numerous protease inhibitors, many of which are members of the serine protease inhibitor (SERPIN) family (13). We have proposed that HDL-mediated delivery of protease inhibitors to plaque may be a novel mechanism for HDL-associated atheroprotection that is independent of HDL cholesterol content (13).

In this manuscript, we describe a novel HDL-targeting protease inhibitor peptide designed to bind predominantly to HDL in plasma and confer elastase-inhibitor activity. Using this peptide, we test the hypothesis that enrichment of HDL-associated elastase inhibitor activity is protective against lesion progression in a mouse model of atherosclerosis.

## Methods

### Mouse housing and diet

Mice were maintained at the University of Kentucky Division of Laboratory Animals Resources in individually vented cages (max. 5 mice per cage) on a 14:10 hour (light/dark) cycle at 22°C (72°F). Teklad Sani-Chip (#7090A, Harlan Teklad) bedding is used in cages and colonies are maintained on a standard rodent diet (#2918 Envigo) with ad libitum access to food and water. All studies were performed in male mice. All animal studies were approved by the University of Kentucky Institutional Animal Care and Use Committee.

### HDL-targeting protease inhibitor peptide design and synthesis

The HTPI peptide was synthesized by a commercial synthesis company at >90% purity confirmed by HPLC and product mass confirmed by mass spectrometry. Elastase inhibitor activity of the peptide was confirmed by in vitro assay (see below). Fluorescent derivatives were generated by N-hydroxysuccinimide chemistry to conjugate fluorophores to the lysine residue situated between the two short polyethylene glycol domains. Labeled peptide was re-purified by reverse-phase chromatography. Alexa 488 labeled HTPI was used for *in vitro* studies and Alexa 647 labeled HTPI was used for *in vivo* studies.

### Elastase activity assays and immunofluorescent detection of neutrophil elastase

*In vitro* elastase activity measurements were performed using elastase activity assay kit (Invitrogen # E12056) with the fluorescent substrate BODIPY-FL-DQ elastin. Elastase activity measurements in whole aorta tissue or tissue sections were performed using the far-red fluorescent elastase substrate Neutrophil Elastase 680 FAST probe (NE680; Perkin Elmer # NEV11169). Immunofluorescent detection of neutrophil elastase was performed on aortic sections from *apoE*^*-/-*^ mice (The Jackson Laboratory strain # 002052). Frozen sections were fixed with fresh 4% paraformaldehyde and neutrophil elastase protein was detected using anti-neutrophil elastase antibody (Abcam #68672).

### Coarse-grain molecular dynamics

Prediction of HTPI interaction with a phospholipid micelle was performed using the Positioning of Proteins in Membranes (PPM) server (version 3.0; University of Michigan) (14, 15). We first generated a molecular structure for the HTPI peptide using Maestro 13.0 (Schrödinger). This structure was then uploaded to the PPM server for positioning in a dodecylphosphocholine (DPC) micelle. Output structure was visualized in Maestro to generate the final model.

### Lipoprotein separation by size-exclusion chromatography

Deidentified human plasma obtained from the Kentucky Blood Center was used for HTPI binding experiments. Alexa 488-labeled HTPI (5 μg) was incubated with human plasma (50 μL) at 37ºC for 1 hour then plasma lipoproteins were separated by fast protein liquid chromatography (FPLC). FPLC was performed on Atka™ Pure instrument with one Superose™ 6 Increase column (Cytiva). Plasma (100 μL) was injected onto the column, and eluted with phosphate-buffered saline at a flow rate of 0.5 mL/min. Fractions (0.5 mL/fraction) were collected in deep-well 96 well plates. Total cholesterol was measured by enzymatic assay (Pointe Scientific) across all collected fractions to visualize lipoprotein distribution. The distribution of HTPI-488 across lipoprotein classes was determined by scanning Alexa 488 fluorescence signal in each fraction.

### Plasma clearance of HTPI in vivo

C57BL/6J (The Jackson Laboratory # 000664) and *Apob*^*TG*^ / *Cetp*^*TG*^ double-transgenic mice were used for *in vivo* studies of plasma clearance of the HTPI peptide. Double transgenic mice were generated by crossing *Apob*^*TG*^ (Taconic #1104) with *Cetp*^*TG*^ (The Jackson Laboratory #3904). The double transgenic mice are continually backcrossed with C57BL/6J (The Jackson Laboratory # 000664). HTPI-488 was administered by retroorbital injection to mice and plasma sampled at several time points minutes after administration. To calculate peptide clearance rates, fluorescence signal from a 2-minute post-administration timepoint was used as the peak plasma concentration and signal from subsequent bleeds calculated as a percentage relative to the peak. Plasma half-life (T_1/2_) was calculated as the time at which plasma HTPI-488 signal reached 50% of the peak.

### Endothelial transcytosis studies

Transcytosis studies were performed using human aortic endothelial cells (HAEC; ATCC # PCS-100-011). Cells were cultured in endothelial basal medium supplemented with an endothelial cell growth kit containing VEGF (ATCC # PCS-100-041). Cells were seeded onto 12 mm Transwell supports at a density of 100,000 cells/well and cultured for 48-72 hours to establish a confluent monolayer, verified by permeability to 40kDa FITC dextran. HTPI-488 was added to the apical chamber in the presence or absence of centrifugally isolated human HDL (50 μg/mL protein). Cells were incubated for 24 hours and media was sampled from the basolateral chamber at several timepoints during the incubation for detection for HTPI-488 fluorescence signal.

### Atherosclerosis studies

Low-density lipoprotein receptor knockout mice (*Ldlr*^*-/-*^) were used to study the impact of HTPI on atherosclerosis. Mice were purchased from The Jackson Laboratory (strain # 002207). Upon initiation of the study, 8-week-old male mice were switched from standard chow to Western diet (WD, 42% kcal from fat + 0.2% cholesterol by weight; TD.88137, Envigo) and maintained on this diet for 12-14 weeks. Studies included three treatment groups: standard chow diet control, WD+saline, and WD+HTPI. *Prevention study*: Saline or HTPI (2.5 mg/kg) were administered by retroorbital injections twice per week for 12 weeks while mice were maintained on WD. *Progression study*: Mice were fed WD for 12 weeks. Starting on week 12, saline or HTPI (2.5 mg/kg) were administered by retroorbital injections three times per week for two weeks while mice were maintained on WD. Aortas were harvested on week 14. For all studies, mice were sacrificed by injectable anesthesia overdose (ketamine 210 mg/kg and xylazine 30 mg/kg) and were perfused with PBS via the left ventricle after severing the right renal artery. Mouse aortas were harvested and fixed in 10% neutral buffered formalin for 24 hours at room temperature, then stored in PBS at 4ºC. Aortas were cleaned thoroughly by removal of periaortic adventitia, stained with Oil Red O, and cut open and pinned flat for en face analysis. Images were taken on Nikon Imaging Software and ImageJ was used to quantify stained plaque areas. Lesion area was calculated (lesion area/total aortic surface area) *100. Aortic roots were embedded in OCT immediately after harvest and kept at -80°C until sectioning. Serial sections (10 μm) were cut and placed on slides (Thermo Fisher Scientific). For immunofluorescence staining, root sections were fixed with 4% paraformaldehyde and probed with antibodies against CD68 (Abcam # ab125212) and smooth muscle actin (Abcam # ab5694). Fluorescence imaging was performed on a Zeiss AxioScan slide scanner.

### Statistical analyses

For statistical comparisons between more than two groups analyses were performed by ANOVA with post-hoc adjustment using methods indicated for each experiment in figure legends. Statistical comparisons were performed using GraphPad Prism software. For all experiments, p values < 0.05 were considered statistically significant.

## Results

### AAPV-CMK inhibits elastase activity in atherosclerotic lesions

To demonstrate the presence of elastase in atherosclerotic lesions, sections of aorta from atherosclerosis prone *Apoe*^*-/-*^ mice were immunostained for detection of neutrophil elastase (NE). NE was detected predominantly in the intimal layer where atherosclerotic lesions were present (yellow arrow, **Figure 1A**). Fresh un-fixed sections incubated with the elastase substrate NE680 resulted in generation of fluorescent signal indicative of active NE in the tissue (**Figure 1B**). Pretreatment of the tissue with a protease inhibitor cocktail (HALT™; Thermo Scientific) prevented proteolytic activity and the generation of fluorescent signal from NE680 (**Figure 1C**). A similar experiment performed on aorta sections from Western diet fed *Ldlr*^*-/-*^ mice demonstrates elastase activity within lesions (**Figure 1D**) that is suppressed by pre-treatment with the peptide inhibitor of elastase Ala-Ala-Pro-Val-chloromethylketone (AAPV-CMK) (**Figure 1E**). NE activity was also detectable in atherosclerosis prone regions of whole aorta tissue from both *Ldlr*^*-/-*^ and *Apoe*^*-/-*^ mice *ex vivo*, and was inhibited by pretreatment with AAPV-CMK (**Figure 1F**).

**Figure 1.**
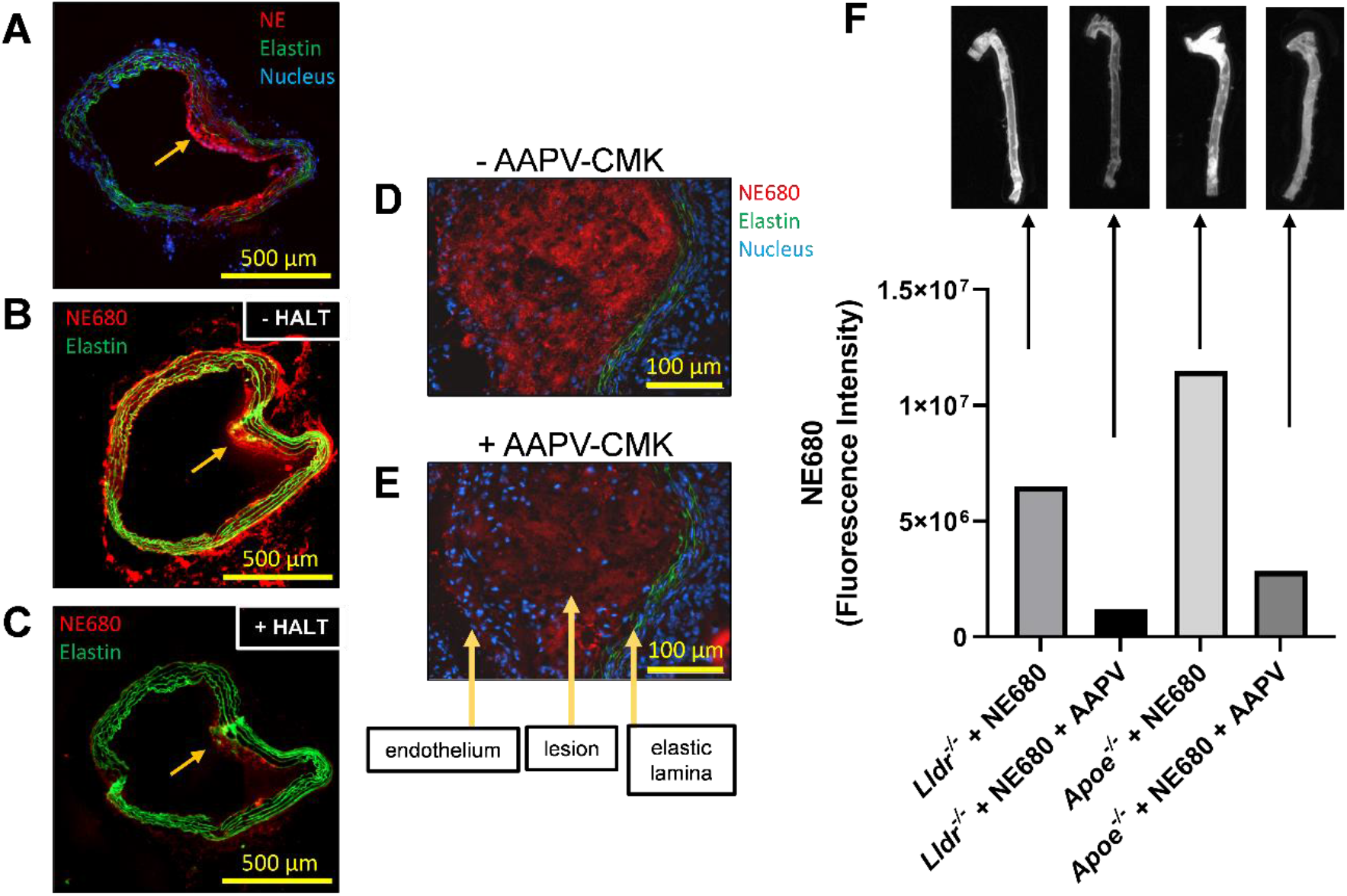
Active neutrophil elastase is present in atherosclerotic lesions and is inhibited by AAPV-CMK. (A) Immunofluorescence detection of neutrophil elastase (NE) in atherosclerotic lesion from aortic section of an *Apoe*^*-/-*^ mouse. Red – neutrophil elastase, green – elastin (autofluorescence), blue – nuclei (DAPI). Arrow indicates location of atherosclerotic lesion. (B-C) Consecutive aortic sections from an *Apoe*^*-/-*^ mouse were treated with the elastase activity probe NE680 in the absence (B) or presence (C) of HALT™ protease inhibitor cocktail (Thermo Scientific). (D-E) Consecutive aortic root sections from *Ldlr*^-/-^ mouse on Western Diet treated with NE680 in the absence (D) or presence (E) of the elastase inhibitor AAPV-CMK. (F) Detection of elastase activity in whole aorta tissue from *Ldlr*^*-/-*^ and *Apoe*^*-/-*^ mice. The elastase activity probe NE680 was used in the absence or presence of the elastase inhibitor AAPV-CMK.

### Design of an HDL-targeting elastase inhibitor

We sought to synthetically increase the elastase inhibitor capacity of HDL by designing a small peptide which could bind to HDL and confer elastase inhibitor activity. To achieve this, we conjugated AAPV-CMK to a molecule of alpha-tocopherol via a short polyethylene glycol spacer (**Figure 2A**). The function of the spacer domain is to increase the distance between AAPV-CMK and the HDL-targeting alpha-tocopherol and to introduce a lysine residue with a primary amine allowing for optional tagging of the peptide by N-hydroxy succinimide chemistry. We refer to this compound as HDL-targeting protease inhibitor (HTPI). We first tested the elastase inhibitor activity of HTPI compared to AAPV-CMK and demonstrate that addition of the spacer and targeting domains does not impact activity (**Figure 2B**). HTPI and AAPV-CMK had elastase inhibitor activity slightly less potent than the human elastase inhibitor protein alpha-1-antitrypsin (AAT). To predict how the HTPI peptide would interact with a lipoprotein particle, we built a molecular model of HTPI and used this as input in to the Position of Proteins in Membranes server to run a coarse-grain molecular dynamics energy minimization with a lipid micelle. This simulation resulted in insertion of the alpha-tocopherol domain into the lipid micelle with the AAPV-CMK domain protruding from the micelle into the solvent (**Figure 2C**).

**Figure 2.**
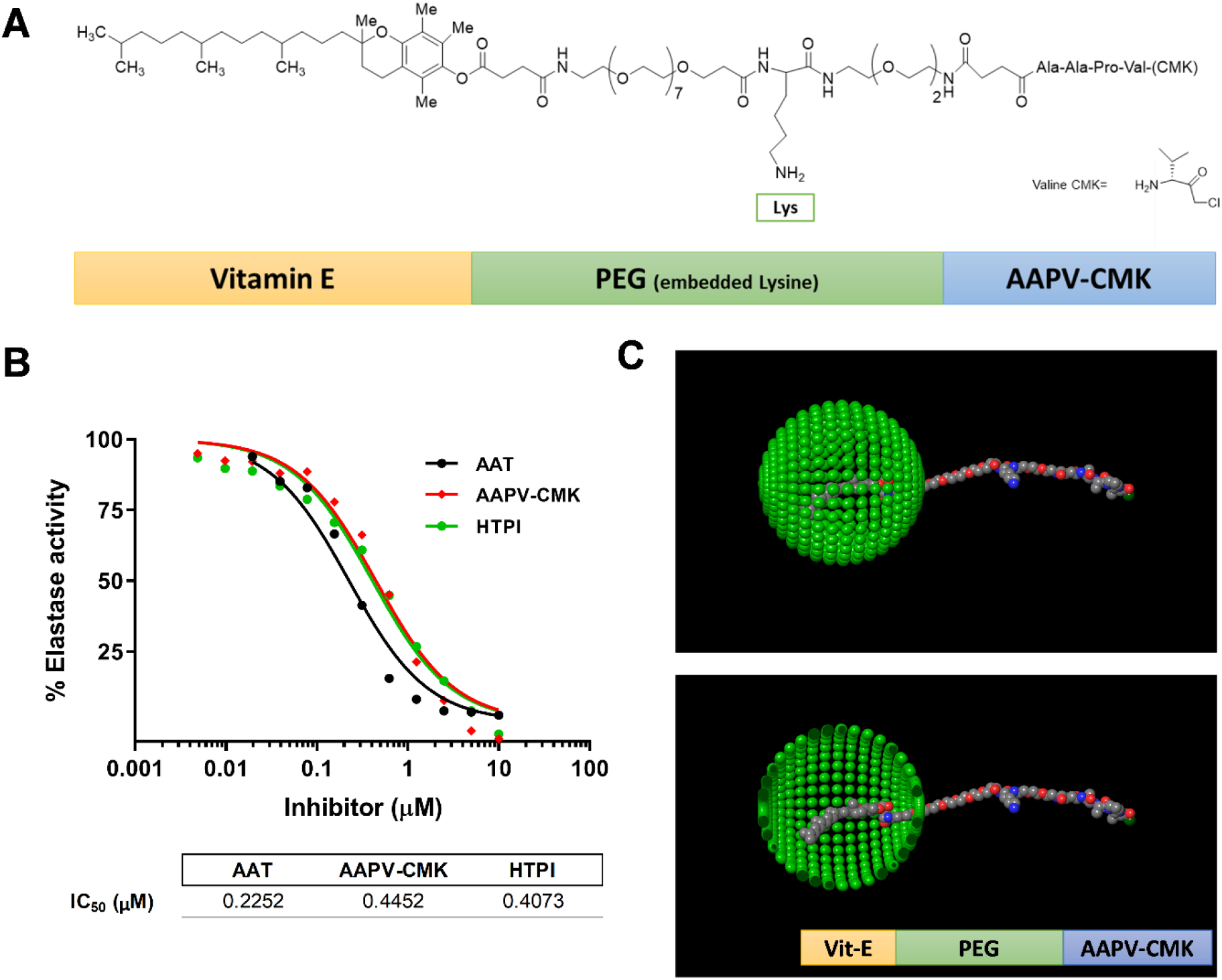
Development of an HDL-targeting protease inhibitor (HTPI) (A) Chemical structure of the HTPI peptide with three domains: HDL-targeting (vitamin E), soluble linker (PEG), and elastase inhibitor (AAPV-CMK). (B) Quantification of elastase inhibitor activity by human alpha-1-antitrypsin protein (AAT), AAPV-CMK, and HTPI. (C) Coarse-grain molecular dynamics simulation of the HTPI interaction with a DPC (green spheres, Dodecylphosphocholine) micelle predicts insertion of the alpha-tocopherol domain into the lipid core. Upper panel displays full micelle spherical structure. Lower panel displays a cutaway of micelle structure to reveal embedded vitamin E domain. Simulations were performed using the Position of Proteins in Membranes (PPM) server (version 3.0; University of Michigan).

### HTPI binds HDL in human plasma *ex vivo* and in mouse plasma *in vivo*

To examine the affinity of HTPI for human lipoproteins, fluorescent HTPI-488 was incubated with human plasma *ex vivo* and then separated by size-exclusion chromatography. HTPI-488 was associated primarily with HDL (68%) and to a lesser extent with other plasma lipoproteins (LDL = 27% and VLDL = 4%) (**Figure 3A**). To examine HTPI binding to lipoproteins in vivo, two mouse strains were used. HTPI also predominantly associated with HDL when injected into *Apob*^*TG*^ / *Cetp*^*TG*^ double-transgenic mice with a humanized lipoprotein profile (**Figure 3B**). The half-life of injected HTPI-488 in these mice was 10.6 hours, a value similar to the half-life of plasma HDL in mice (**Figure 3C**). Plasma clearance studies performed in C57BL6-J mice demonstrate a similar half-life for HTPI-488 (T_1/2_=9.4 hrs.) and much shorter half-life’s for the AAPV inhibitor without the HDL-targeting motif (AAPV-488; T_1/2_=1.4 hrs.) or the free Alexa-488 dye (T_1/2_=0.3 hrs.) (**Figure 3D**). Plasma collected 1-hour post-injection with either saline or HTPI was separated by FPLC and elastase inhibitor activity was measured across fractions. Mice injected with HTPI had increased elastase-inhibitor capacity in HDL-containing fractions (**Figure 3E**).

**Figure 3.**
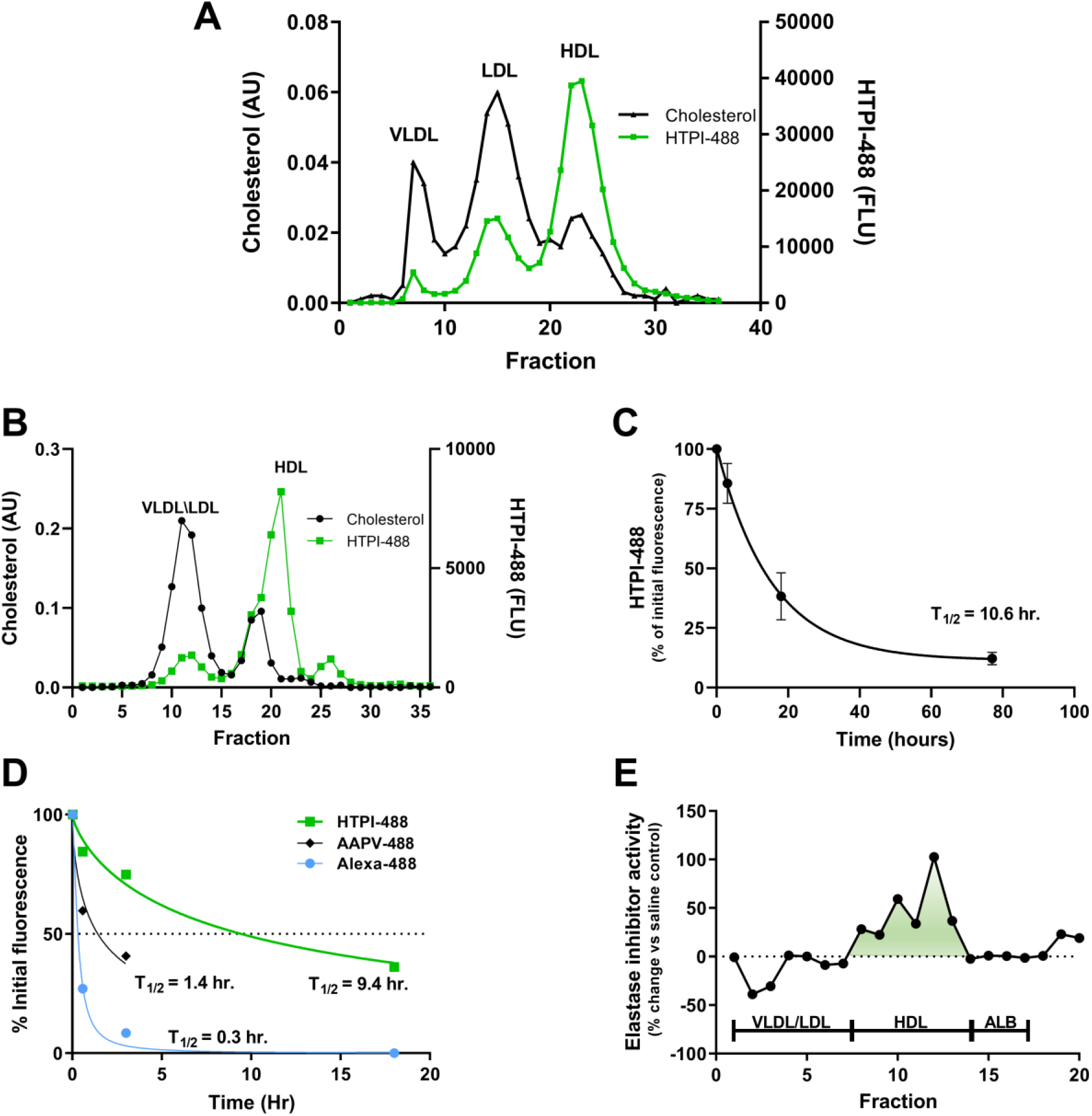
HTPI binds plasma HDL and confers protease inhibitor activity. (A) HTPI binding to lipoproteins in human plasma. Alexa 488 labeled HTPI (HTPI-488) was incubated with human plasma for 1 hour at 37ºC *ex vivo* and plasma lipoproteins were then separated by size-exclusion chromatography using Superose 6 columns. Cholesterol content of fractions was measured by enzymatic assay to determine the distribution of the major lipoprotein classes. Fractions were scanned for fluorescence to assess HTPI distribution among lipoprotein fractions. (B) HTPI-488 was injected to *Apob*^*TG*^ / *Cetp*^*TG*^ double transgenic mice (n=3 mice) and plasma collected after 1 hour was used for size-exclusion chromatography to determine lipoprotein distribution of HTPI. (C) Plasma clearance of HTPI-488 in *Apob*^*TG*^ / *Cetp*^*TG*^ double transgenic mice. (D) Plasma clearance of Alexa-488 (free dye), HTPI-488, and AAPV-488 (control peptide with no HDL-targeting domain) in C57BL6J mice. (E) Elastase inhibitor activity measured in size-exclusion chromatography fractions of plasma collected from mice 1 hour after injection with saline or HTPI. The change in elastase inhibitor activity in lipoprotein fractions relative to saline control is presented. Increased elastase inhibitor activity is observed in HDL fractions of mice treated with HTPI. AU – absorbance units; FLU – fluorescence units; ALB – albumin.

### HDL facilitates endothelial transcytosis of HTPI *in vitro* and systemic administration *in vivo* results in detection of HTPI in plaque

To gain access to extravascular tissues, the HTPI peptide on HDL would need to cross the vascular endothelium. Endothelial transcytosis was measured *in vitro* across Human Aortic Endothelial Cells (HAEC) cultured on membrane supports. Addition of HTPI-488 to the apical chamber resulted in very little transcytosis of the peptide to the basolateral chamber (**Figure 4A**). However, addition of HDL significantly increased the rate of HTPI-488 transport to the basolateral compartment (13.4 vs. 151.4 FL/hr, p<0.0001). To examine HTPI uptake in atherosclerotic lesions *in vivo*, HTPI-647 was administered to *Apoe*^*-/-*^ mice by retroorbital injection. HTPI-647 signal was detected via fluorescence scanning of whole aortas ex vivo and accumulated in regions that are prone to atherosclerosis, particularly in the aortic arch (**Figure 4B**). Fluorescence microscopy of aortic sections revealed accumulation of HTPI-647 within atherosclerotic lesions (**Figure 4C**).

**Figure 4.**
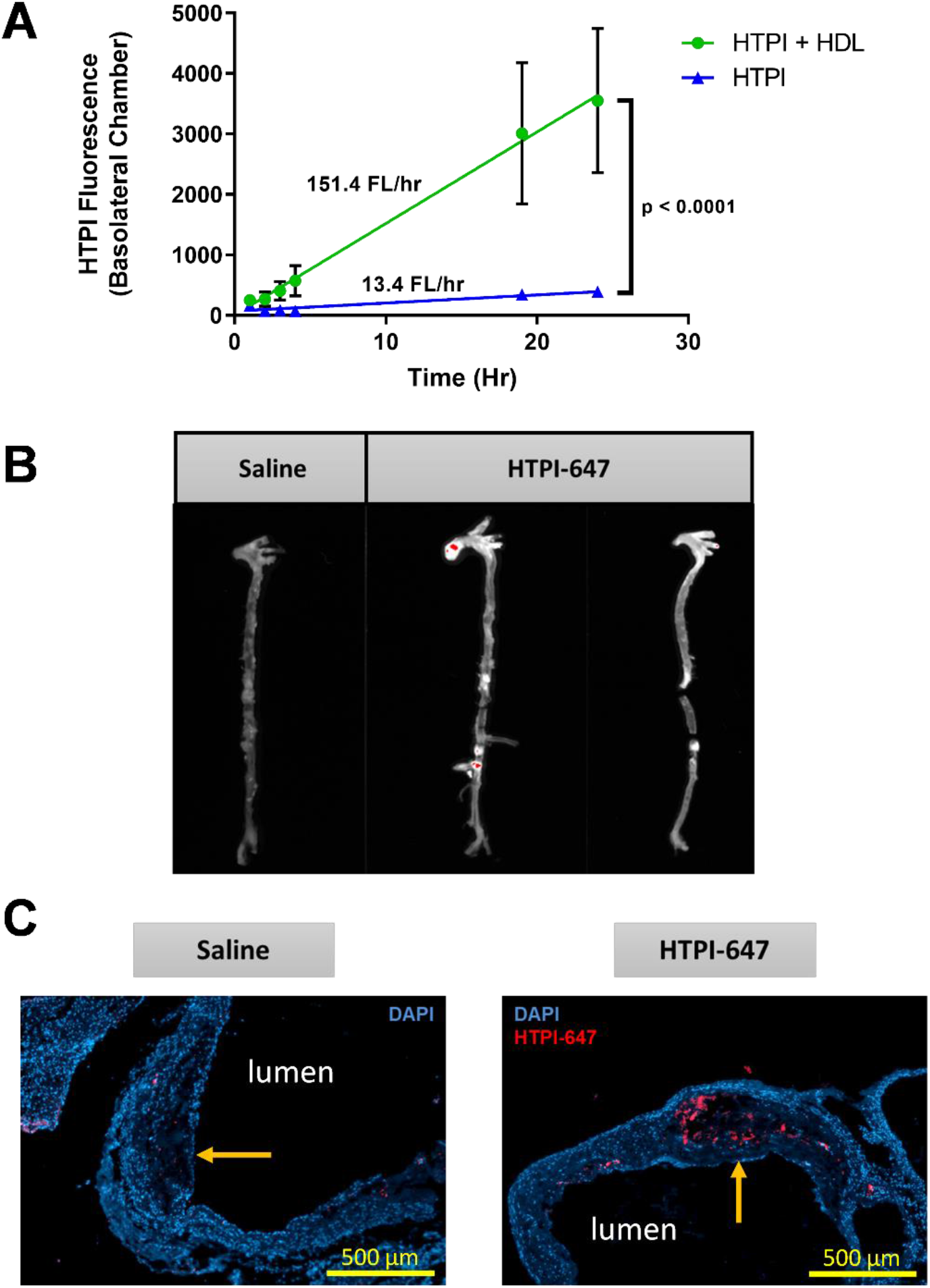
HDL facilitates endothelial transcytosis of HTPI *in vitro* and systemic administration *in vivo* results in detection of HTPI in plaque. (A) Endothelial transcytosis of fluorescent HTPI. Human aortic endothelial cells were cultured on membrane supports and HTPI-488 was added to the apical chamber in the absence or presence of HDL (50 μg/mL protein). Accumulation of HTPI-488 fluorescence in the basolateral chamber was monitored over time (n=3/group). (B) Detection of HTPI-647 in whole aortic tissue of *Apoe*^*-/-*^ mice after retroorbital injection. (C) Fluorescence microscopy of aortic sections from *Apoe*^*-/-*^ mice injected with saline or HTPI-647. Plaque (orange arrows) of mice who received HTPI-647 show accumulation of fluorescence signal. Statistical comparisons in panel (A) performed by least squares regression and comparison of slopes by the extra sum-of-squares F test.

### HTPI reduces atherosclerosis lesion area in *Ldlr*^*-/-*^ mice

The impact of enhanced HDL-associated elastase inhibitor activity on atherosclerosis was evaluated in two studies using *Ldlr*^-/-^ mice (**Figure 5A**). For the Prevention Study, mice were fed Western diet (WD) for 12 weeks while receiving twice weekly retroorbital injections of saline or HTPI (2.5 mg/kg). For the Progression Study, *Ldlr*^-/-^ mice were fed a Western diet for 12 weeks, then administered retroorbital injections of saline or HTPI (2.5 mg/kg) three times per week for two weeks while maintained on Western diet. Atherosclerosis lesion area was quantified by en face analysis of oil red O stained aortas (**Figure 5B**). In both studies, mice fed WD developed significant atherosclerosis compared to standard diet fed mice. In the Prevention Study, HTPI-treated mice had 21% smaller plaque area compared to saline-treated mice (10.7% vs 13.6%; q<0.05). In the Progression Study, HTPI-treated mice had 28% smaller plaque area compared to saline-treated mice (13.4% vs 18.6%; q<0.01). Comparison between the two studies reveals that HTPI-treated mice in the Progression Study have similar plaque area to saline-treated mice in the Prevention Study (**Figure 5B**, horizontal dotted line), revealing that the two-week HTPI treatment in the Progression Study completely prevented the increase in lesion area observed in the saline-treated group. Plaque morphology was examined in aortic root sections from the Progression Study. Oil red O staining of sections revealed reduction in lipid content as observed by en face aorta analysis (**Figure 5C**). Immunostaining of aortic root sections for the macrophage marker Cd68 indicated a 28% reduction in lesion macrophage content in the HTPI-treated mice (**Figure 5C**,**D**). Immunostaining for smooth muscle actin (Acta2) revealed a possible effect of HTPI on the luminal surface of plaque, with thinner and less-fragmented actin strands in HTPI-treated mice. This may be indicative of a possible stabilizing effect of HTPI on the fibrous cap.

**Figure 5.**
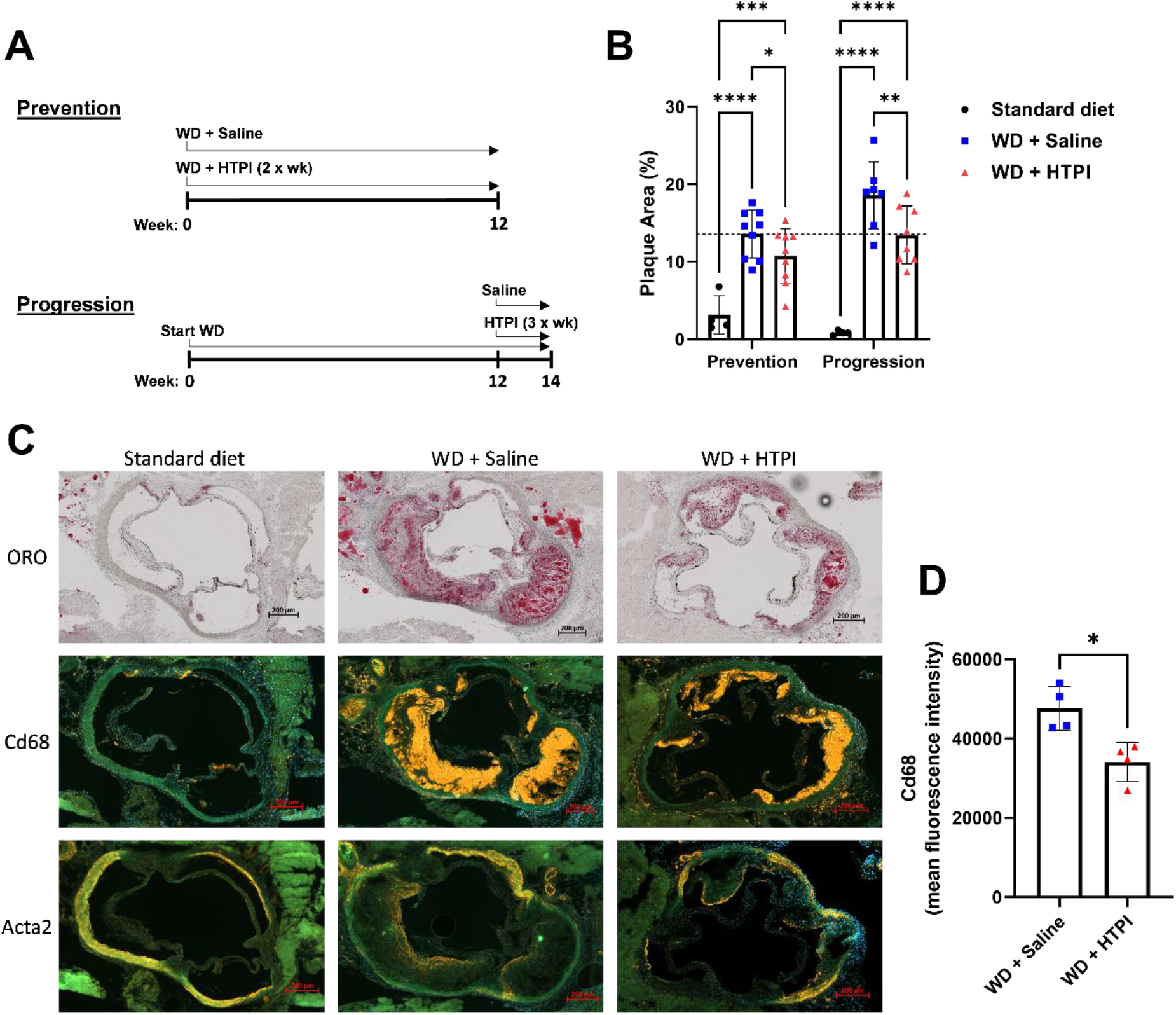
HTPI reduces atherosclerosis lesion area in *Ldlr*^*-/-*^ mice. (A) Study designs for atherosclerosis Prevention and Progression using *Ldlr*^*-/-*^ mice fed Western diet. HTPI dose for both studies was 2.5 mg/kg body weight. (B). Plaque burden was quantified by oil-red-o staining of whole aorta tissue and en face imaging was used for quantification of plaque area as a percentage of total area. (C) Plaque morphology of aortic root sections from the Prevention study: Oil Red O (ORO) staining, Cd68 and smooth muscle actin (Acta2) immunostaining. (D) Cd68 fluorescence intensity measured from aortic root sections from the Prevention study. Statistical comparisons: (B) performed by two-way ANOVA with multiple comparisons correction by the method of Benjamini, Krieger, and Yekutieli. *q>0.05; **q>0.01; ***q>0.001; ****q>0.0001. (D) Unpaired two-tailed t test. *p>0.05.

## Discussion

A variety of proteases are secreted within atherosclerotic lesions and these are generally believed to contribute to progression and destabilization of the plaque. Since endogenous HDL carry an array of protease inhibitor proteins, we hypothesized that enrichment of protease inhibitor activity on HDL may prevent lesion progression and promote plaque stability. In this manuscript, we present a novel approach for targeted functional enhancement of lipoproteins using a vitamin E conjugated peptide that binds predominantly to HDL in plasma. Using this strategy, we designed an HDL-targeting elastase-inhibitor peptide to enrich the elastase-inhibitor activity on HDL and then examined the impact of administration of this peptide on atherosclerosis lesion progression in a mouse model of atherosclerosis.

Elastase was chosen as the target protease for this first generation of the HTPI peptide. Elastase is present within atherosclerotic lesions in humans and mice (8). Within plaque, it is secreted by both neutrophils and macrophages (8), however, the contribution of this specific protease to atherosclerosis progression or lesion stability is not well understood. Genetic deficiency of elastase results in protection against atherosclerosis in *Apoe*^*-/-*^ mice (16). Furthermore, systemic administration of a small molecule elastase inhibitor has been demonstrated to protect against atherosclerosis in *Apoe*^*-/-*^ mice (16). Our HTPI peptide offers some advantages over systemic administration of a small molecule inhibitor. Targeting of HTPI to HDL results in reduced clearance and a longer half-life, which could result in lower dosing requirements and less frequent administration. Additionally, the plaque-targeting properties of HTPI-HDL may reduce off-target effects of systemic elastase inhibition. It is of note that, the physiological inhibitor of elastase, alpha-1-antitrypsin, has been detected on both human and mouse HDL in numerous proteomics studies (11, 12, 17). Although the functional and structural basis for this interaction is not known, we recently published that there may be a functional benefit from the interaction between AAT and HDL, specifically an increased anti-inflammatory potential of AAT in the presence of HDL (18). HDL accumulate within atherosclerotic lesions where they can efflux cholesterol from macrophage foam cells and promote reverse cholesterol transport (9, 10). We hypothesized that AAT may confer elastase-inhibitor potential on to HDL and that this may be a novel mechanism by which HDL can be atheroprotective (13).

The current study sought to mimic the function of AAT protein with a small peptide inhibitor of elastase, AAPV-CMK, and to test the impact of enrichment of this activity on HDL in atherosclerosis development and progression. This strategy offered some functional advantages over the whole AAT protein. One key advantage is the capacity for enrichment of elastase-inhibitor activity on HDL. We have previously estimated that human HDL can only carry 1-2 molecules of AAT per particle (18). The HTPI peptide is approximately 1/30^th^ the size of AAT protein, thus there is a potential for greater loading of HDL with HTPI, on a per-particle basis although this is difficult to measure due to a lack of methods for absolute quantification of the synthetic peptide.

By measuring elastase activity in mouse atherosclerosis lesions, we confirmed previous studies detecting significant elastase activity in aortic roots and in whole aorta tissue. We next demonstrated that the AAPV-CMK peptide can inhibit the elastase activity detected within lesions, supporting its suitability as a tool to influence proteolytic activity within plaque. The HTPI peptide design joins AAPV-CMK to vitamin E (alpha tocopherol) via a polyethylene glycol linker that serves both as a physical spacer and a means to improve aqueous solubility of the peptide. Importantly, we demonstrate that conjugation of AAPV-CMK to vitamin E does not result in any loss of activity compared to unconjugated AAPV-CMK. Furthermore, the HTPI peptide had only modestly lower elastase-inhibitor activity, on a molar basis, compared to AAT protein. Molecular dynamics simulation of HTPI binding to a lipid micelle supports our intended design for interaction of HTPI with lipoproteins via the vitamin E domain and extension of the protease inhibitor domain from the lipoprotein particle making it available for interaction with soluble proteases.

The majority of HTPI was bound to HDL when incubated with human plasma, where significant amounts of apoB-containing lipoproteins are present. In mouse plasma, the amount of HTPI on apoB lipoproteins was likely lower because mice have considerably lower concentrations of VLDL and LDL. However, in apoB/CETP double transgenic mice, that have a “humanized” lipoprotein profile, the majority of HTPI was bound to HDL. The plasma half-life of HTPI in these studies was consistent with the half-life of mouse HDL (19). The AAPV-CMK peptide without the HDL-targeting domain had about a 7-fold shorter half-life and was rapidly detected in urine (within minutes) likely due to renal clearance of the <1 kDa free peptide. The lack of absolute specificity for HDL is a minor limitation of this study, since the targeting of HTPI to plaque cannot be completely attributed to HDL and there may be a contribution of HTPI delivery to plaque by peptide bound to apoB-containing lipoproteins. This may also represent an effective route of delivery of compounds to lesions and does not diminish the significance of the finding that enriching elastase inhibitor activity on lipoproteins significantly prevented increases in lesion size.

Accumulation of HTPI within atherosclerotic lesions is consistent with the accumulation of HDL within plaque that has previously been reported (20). We hypothesized that HTPI is co-transported into the lesion during endothelial transcytosis of HDL. Endothelial transcytosis of HDL was demonstrated by Von Eckardstein et. al. to be mediated via SCARB1 and ABCG1 for spherical HDL and via ABCA1 for discoidal HDL (21, 22). Consistent with our hypothesis, our *in vitro* data demonstrate that endothelial transcytosis of HTPI is HDL-dependent. Furthermore, our *in vivo* data demonstrate localization of HTPI in atherosclerosis-prone regions of the vasculature. Collectively, these data indicate that our peptide-based approach is an effective way to target atherosclerotic lesions.

Administration of HTPI three times weekly for two weeks to mice with established atherosclerosis prevented additional lipid accumulation and expansion of plaque area, but did not result in plaque regression. These data suggest that local inhibition of elastase within the lesion can prevent plaque progression, but may not necessarily facilitate the processes involved in lesion reduction in the timeline studied here. It is not yet clear what the exact mechanism underlying this effect are, but there are several possiblities. It may be that inhibition of the intralesional elastase activity reduces endothelial activation and subsequent recruitment of inflammatory cells into the plaque. This is supported to some extent by immunostaining of aortic root sections, which revealed lower Cd68 content in lesions of HTPI-treated mice. Another possible mechanism is that HTPI administration results in reduced uptake of apoB-containing lipoproteins by macrophage foam cells. Interestingly, elastase-treated LDL display increased uptake by macrophages (23, 24). This would suggest that HTPI-mediated inactivation of elastase within plaque could reduce macrophage uptake of LDL and subsequently reduce foam cell formation. Endothelial transcytosis of apoB-containing lipoproteins into the plaque could also be impacted by HTPI. This could be mediated through either direct or indirect effects of elastase on vascular endothelium expression of LDL transporters, such as SCARB1 (25). Future studies will address these possibilities.

## Conclusions

In summary, our data demonstrate that (a) the HTPI peptide binds HDL and enhances HDL-associated protease inhibitor activity *in vivo*, (b) HTPI accumulates within atherosclerotic lesions in mice, and (c) short-term administration of HTPI prevents lesion progression in a mouse model of atherosclerosis. The presented data support the hypothesis that HDL-associated anti-elastase activity could contribute to the atheroprotective functions of HDL and highlight the potential utility of enrichment of HDL with anti-protease activity for stabilization of high-risk atherosclerotic lesions. A similar peptide-based approach targeting other proteases could be achieved by swapping out the protease inhibitor domain with other small peptide inhibitors targeting different proteolytic enzymes. This could provide opportunities to target a broad range of physiological targets and pathways with known involvement in atherosclerosis or other pathologic conditions.

## References

1. Libby, P., J. E. Buring, L. Badimon, G. K. Hansson, J. Deanfield, M. S. Bittencourt, L. Tokgözoğlu, and E. F. Lewis. 2019. Atherosclerosis. Nature Reviews Disease Primers 5: 56.

2. Ference, B. A., H. N. Ginsberg, I. Graham, K. K. Ray, C. J. Packard, E. Bruckert, R. A. Hegele, R. M. Krauss, F. J. Raal, H. Schunkert, G. F. Watts, J. Borén, S. Fazio, J. D. Horton, L. Masana, S. J. Nicholls, B. G. Nordestgaard, B. v. d. Sluis, M.-R. Taskinen, L. Tokgözoğlu, U. Landmesser, U. Laufs, O. Wiklund, J. K. Stock, M. J. Chapman, and A. L. Catapano. 2017. Low-density lipoproteins cause atherosclerotic cardiovascular disease. 1. Evidence from genetic, epidemiologic, and clinical studies. A consensus statement from the European Atherosclerosis Society Consensus Panel. European Heart Journal 38: 2459–2472.

3. Goldstein, J. L., Y. K. Ho, S. K. Basu, and M. S. Brown. 1979. Binding site on macrophages that mediates uptake and degradation of acetylated low density lipoprotein, producing massive cholesterol deposition. Proc National Acad Sci 76: 333–337.

4. Bobryshev, Y. V. 2006. Monocyte recruitment and foam cell formation in atherosclerosis. Micron 37: 208–222.

5. Soehnlein, O., L. Lindbom, and C. Weber. 2009. Mechanisms underlying neutrophil-mediated monocyte recruitment. Blood 114: 4613–4623.

6. Garcia-Touchard, A., T. D. Henry, G. Sangiorgi, L. G. Spagnoli, A. Mauriello, C. Conover, and R. S. Schwartz. 2005. Extracellular proteases in atherosclerosis and restenosis. Arteriosclerosis, thrombosis, and vascular biology 25: 1119–1127.

7. Slack, M. A., and S. M. Gordon. 2019. Protease Activity in Vascular Disease. Arterioscler Thromb Vasc Biol 39: e210–e218.

8. Dollery, C. M., C. A. Owen, G. K. Sukhova, A. Krettek, S. D. Shapiro, and P. Libby. 2003. Neutrophil Elastase in Human Atherosclerotic Plaques Production by Macrophages. Circulation 107: 2829–2836.

9. Rohatgi, A., A. Khera, J. D. Berry, E. G. Givens, C. R. Ayers, K. E. Wedin, I. J. Neeland, I. S. Yuhanna, D. R. Rader, J. A. de Lemos, and P. W. Shaul. 2014. HDL Cholesterol Efflux Capacity and Incident Cardiovascular Events. The New England Journal of Medicine 371: 2383–2393.

10. Ouimet, M., T. J. Barrett, and E. A. Fisher. 2019. HDL and Reverse Cholesterol Transport. Circulation Research 124: 1505–1518.

11. Vaisar, T., S. Pennathur, P. S. Green, S. A. Gharib, A. N. Hoofnagle, M. C. Cheung, J. Byun, S. Vuletic, S. Kassim, P. Singh, H. Chea, R. H. Knopp, J. Brunzell, R. Geary, A. Chait, X.-Q. Zhao, K. Elkon, S. Marcovina, P. Ridker, J. F. Oram, and J. W. Heinecke. 2007. Shotgun proteomics implicates protease inhibition and complement activation in the antiinflammatory properties of HDL. Journal of Clinical Investigation 117: 746–756.

12. Gordon, S. M., J. Deng, J. L. Lu, and S. W. Davidson. 2010. Proteomic Characterization of Human Plasma High Density Lipoprotein Fractionated by Gel Filtration Chromatography. Journal of Proteome Research 9: 5239–5249.

13. Gordon, S. M., and A. T. Remaley. 2017. High density lipoproteins are modulators of protease activity: Implications in inflammation, complement activation, and atherothrombosis. Atherosclerosis 259: 104–113.

14. Lomize, M. A., I. D. Pogozheva, H. Joo, H. I. Mosberg, and A. L. Lomize. 2012. OPM database and PPM web server: resources for positioning of proteins in membranes. Nucleic Acids Res 40: D370–D376.

15. Lomize, A. L., S. C. Todd, and I. D. Pogozheva. 2022. Spatial arrangement of proteins in planar and curved membranes by PPM 3.0. Protein Sci 31: 209–220.

16. Wen, G., W. An, J. Chen, E. M. Maguire, Q. Chen, F. Yang, S. W. A. Pearce, M. Kyriakides, L. Zhang, S. Ye, S. Nourshargh, and Q. Xiao. 2018. Genetic and Pharmacologic Inhibition of the Neutrophil Elastase Inhibits Experimental Atherosclerosis. Journal of the American Heart Association 7.

17. Davidson, W. S., A. S. Shah, H. Sexmith, and S. M. Gordon. 2021. The HDL Proteome Watch: Compilation of studies leads to new insights on HDL function. Biochimica et Biophysica Acta (BBA) - Molecular and Cell Biology of Lipids 1867: 159072.

18. Gordon, S. M., B. McKenzie, G. Kemeh, M. Sampson, S. Perl, N. S. Young, M. B. Fessler, and A. T. Remaley. 2015. Rosuvastatin Alters the Proteome of High Density Lipoproteins: Generation of alpha-1-antitrypsin Enriched Particles with Anti-inflammatory Properties. Molecular & Cellular Proteomics 14: 3247–3257.

19. Kuai, R., D. Li, Y. E. Chen, J. J. Moon, and A. Schwendeman. 2016. High-Density Lipoproteins: Nature’s Multifunctional Nanoparticles. Acs Nano 10: 3015–3041.

20. Huang, Y., J. A. DiDonato, B. S. Levison, D. Schmitt, L. Li, Y. Wu, J. Buffa, T. Kim, G. S. Gerstenecker, X. Gu, C. S. Kadiyala, Z. Wang, M. K. Culley, J. E. Hazen, A. J. DiDonato, X. Fu, S. Z. Berisha, D. Peng, T. T. Nguyen, S. Liang, C.-C. Chuang, L. Cho, E. F. Plow, P. L. Fox, V. Gogonea, W. W. H. Tang, J. S. Parks, E. A. Fisher, J. D. Smith, and S. L. Hazen. 2014. An abundant dysfunctional apolipoprotein A1 in human atheroma. Nature Medicine 20: 193–203.

21. Cavelier, C., L. Rohrer, and A. von Eckardstein. 2006. ATP-Binding cassette transporter A1 modulates apolipoprotein A-I transcytosis through aortic endothelial cells. Circ Res 99: 1060–1066.

22. Rohrer, L., P. M. Ohnsorg, M. Lehner, F. Landolt, F. Rinninger, and A. von Eckardstein. 2009. High-density lipoprotein transport through aortic endothelial cells involves scavenger receptor BI and ATP-binding cassette transporter G1. Circulation research 104: 1142–1150.

23. Polacek, D., R. E. Byrne, G. M. Fless, and A. M. Scanu. 1986. In vitro proteolysis of human plasma low density lipoproteins by an elastase released from human blood polymorphonuclear cells. Journal of Biological Chemistry 261: 2057–2063.

24. Polacek, D., R. E. Byrne, and A. M. Scanu. 1988. Modification of low density lipoproteins by polymorphonuclear cell elastase leads to enhanced uptake by human monocyte-derived macrophages via the low density lipoprotein receptor pathway. Journal of Lipid Research 29: 797–808.

25. Huang, L., K. L. Chambliss, X. Gao, I. S. Yuhanna, E. Behling-Kelly, S. Bergaya, M. Ahmed, P. Michaely, K. Luby-Phelps, A. Darehshouri, L. Xu, E. A. Fisher, W.-P. Ge, C. Mineo, and P. W. Shaul. 2019. SR-B1 drives endothelial cell LDL transcytosis via DOCK4 to promote atherosclerosis. Nature 569: 565–569.

